# A Wirelessly Powered Implantable CMOS Neural Recording Sensor Array using Pulse-based Neural Amplifier

**DOI:** 10.1101/809509

**Authors:** Geoffrey Mulberry, Kevin A. White, Brian N. Kim

## Abstract

The most important organ in the human body is unquestionably the brain. Yet, despite its importance, it is one of the least well understood organs. One reason for this lack of understanding of the brain is the lack of data available to researchers from in vivo studies. Historically, collecting measurements from the brain has been difficult due to the high risk to the patient. Recently technology has been developed to allow electrical measurements to be taken from the brain directly, however most systems involve non-permanent sensors because of the requirement for transcranial wiring for power and data. Developments in the field of CMOS circuit design, wireless power transmission, and wireless data transmission have enabled the creation of implantable neural recording devices as a combination of these technologies. The implant designed in this paper is ∼15 mm in diameter and 2 mm at its thickest point on a flexible polyamide PCB. The flexible nature of the implant allows for the implant to conform to the surface of the brain. The implant requires no transcranial orifice since it is powered wirelessly and transmits data wirelessly via Bluetooth low energy. The CMOS neural amplifier chip on the implant utilizes an enhanced form of delta modulation to remove the requirement for large ADCs to be present on the die, saving space and enabling 1024 amplifiers and electrodes to be present on the chip. The implant is capable of measuring, modulating, and wirelessly transmitting a millivolt order signal to a PC for demodulation and analysis.

## I. INTRODUCTION

NEURAL recording implants offer great potential for studying the human brain in ways that were impossible just a few decades ago. This has inspired much research in the field, including three separate technologies: high-throughput complementary metal-oxide semiconductor (CMOS) array chips, high performance microelectrodes, and wireless power transmission (WPT) methods[1]. Some have spent research effort on CMOS sensor arrays in an effort to integrate a large number of high-performance amplifiers onto a single chip [2]–[7]. Others have focused on fabricating high quality electrodes designed specifically for probing into the brain [8]–[11]. Many have also powered these type of CMOS devices wirelessly [12]–[15]. Combining these technologies will enable studies of the brain to take place at a scale and density that was not before possible.

This study focuses on a device combining two of these technologies, a highly integrated CMOS neural recording chip, with wireless power transfer. CMOS amplifier arrays can be designed for any high throughput applications where a large number of parallel electrical measurements can be made. The CMOS device fabricated for this study is optimized for measuring neural signals while also integrating a high electrode count of 1024. To achieve this high electrode count, while still allowing for a wireless data transmission, this CMOS device utilizes a pulse-based neural amplifier. Utilizing enhanced delta modulation (EDM) enables the high amplifier count by placing a modulator in each array element, making the outputs from each amplifier digital, rather than analog. This removes the requirement for one of the largest features traditionally found on CMOS amplifier arrays, the analog-to-digital converter (ADC). In fact, designing compact ADCs is a popular research topic [16]–[18]. In addition to removing the ADCs, placing a modulator in each array element has added benefits since digital signals are better suited to multiplexing and less susceptible to signal-related problems such as noise than analog signals. Additionally, the signals being digital immediately out of the amplifiers lends itself to high scalability, since additional multiplexing requires little circuitry and needs only a higher bandwidth communication channel. A comparable analog system would require either faster (and therefore higher performance) ADCs, or additional ADCs to scale to higher array element counts. Both of these solutions would consume significant CMOS die area and power, effectively providing a limit for the number of amplifiers that can be integrated onto a single chip of the same size.

In this paper, the presented CMOS chip is mounted to a polyimide flexible printed circuit board (PCB) which contains additional electronics to facilitate a complete neural recording implant. This flexible substrate enables the system’s wireless power transfer by using spiral traces as an inductive coil. It also holds a system-on-chip (SOC) that operates the CMOS chip and sends data wirelessly via Bluetooth low energy (BLE). The flexible nature of the thin polyimide-based substrate enables the implant to bend, allowing for the device to conform to the surface of the brain.

This paper discusses the design of the flexible neural implant and tests of its functionality. Section II describes the overall design of the device, describing the flexible substrate. In Section III, the circuitry and design of the coils for the wireless power transfer link are detailed. Section IV focuses on the neural recording amplifier design and the operation of the pulsed-based amplifier circuitry. To demonstrate the functionality of the device, Section V presents results from the fabrication of the flexible implant and measurements of the wireless power transfer system, the neural amplifiers, the enhanced delta modulators, and the Bluetooth data transfer system.

## II. FLEXIBLE NEURAL RECORDING IMPLANT

The flexible neural recording implant is a compact disk shaped implant that is wirelessly powered and thin enough to be implanted directly on the surface of the brain under the skull. An overview of the system is shown in Fig. 1. The implant is small, only 15.4 mm in diameter by 2.0 mm at its thickest. The small size allows it to be placed on the brain under the skull. The flexible neural recording implant is fully wireless and does not require any wiring or battery to operate. Power is provided by an external coil, and data is transmitted wirelessly via BLE to a receiving device such as a laptop computer.

**Fig. 1.**
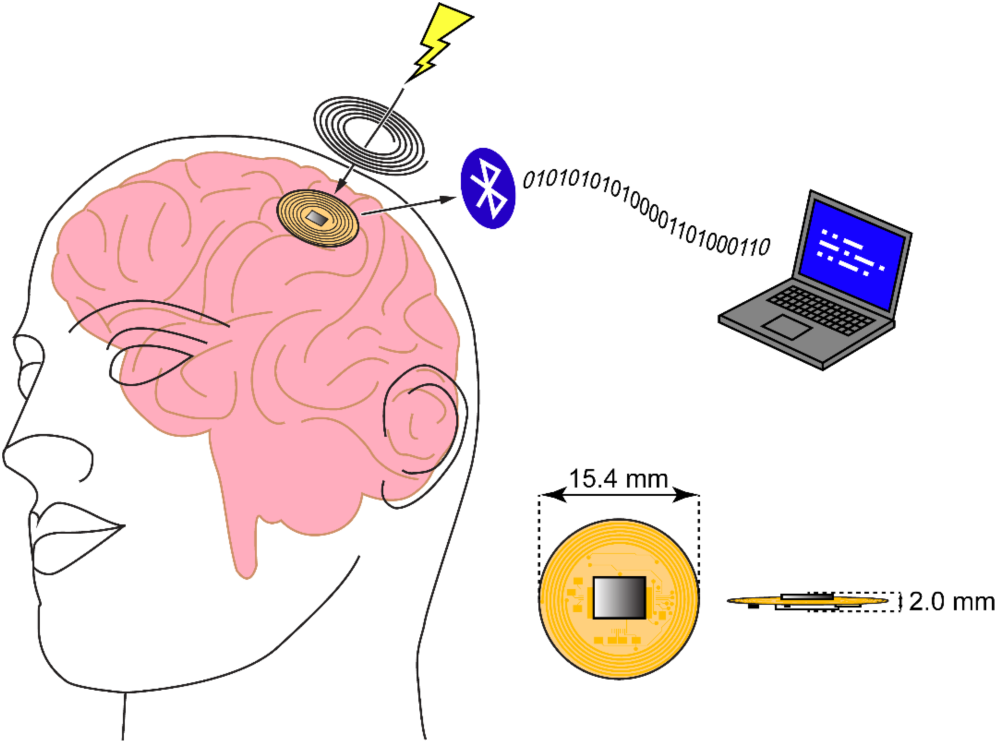
The implantable sensor is a small 15.4 mm 2.0 mm. Its small size is achieved by the use of wireless power transfer provided by an external coil and the flexible substrate. The device transmits measurement data via a low energy Bluetooth link to a receiving device such as a laptop computer for processing.

### A. Flexible implant substrate

The core of the implant is the 15.4 mm diameter by 0.2 mm polyimide flexible substrate. Within this small space, the flexible implant contains electronics for the wireless power supply, the CMOS neural chip, and the SOC. A block diagram for the flexible neural interface is shown in Fig. 2. The wireless power supply consists of a power coil that is implemented using spiral copper traces around the outer annulus of the substrate. This coil is tuned to resonance via a parallel capacitor and fed into a full-wave bridge rectifier. The output of the rectifier is regulated using a 3.3 V regulator that powers the electronics in the neural interface. Signals are recorded via the EDM amplifier array integrated into the CMOS neural chip and sent to the SOC, a CC2640 (SimpleLink ultra-low power wireless MCU for Bluetooth low energy) by Texas Instruments. The SOC contains an ARM microcontroller that generates the required timing signals to operate the CMOS neural chip and processes and packages the data it receives to send via Bluetooth Low-energy. The SOC also contains a built-in BLE transceiver which is used to drive the external chip antenna through a balun.

**Fig. 2.**
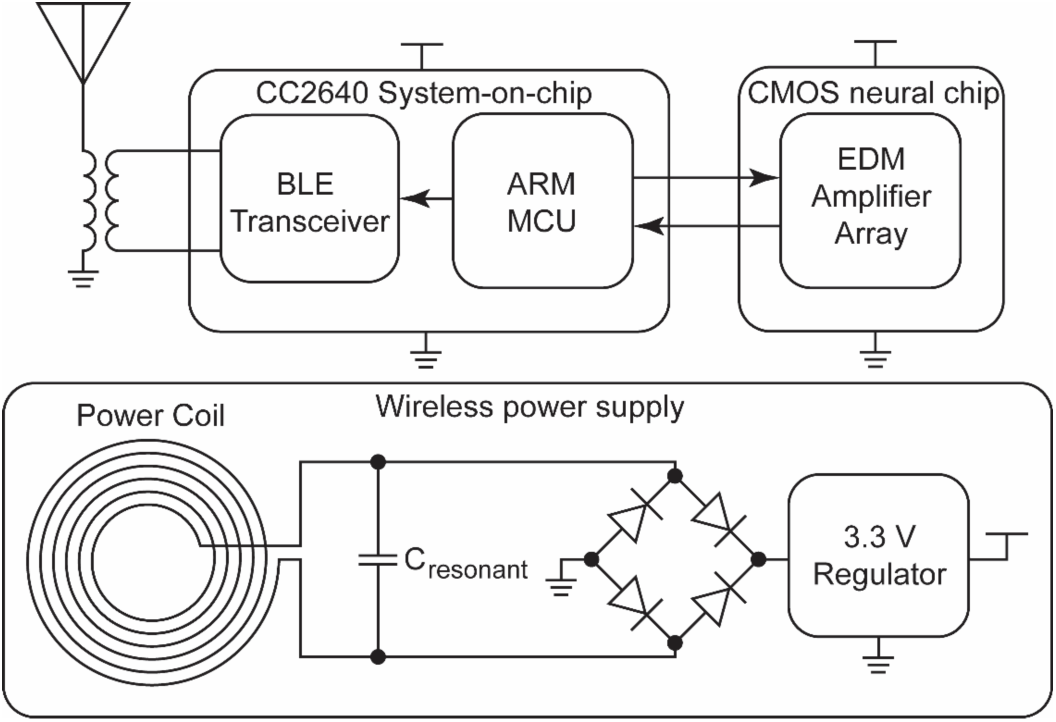
Block diagram of the implantable device. The three main parts are the CC2640 system-on-chip, responsible for controlling the CMOS chip via the ARM microcontroller, as well as driving the antenna via the Bluetooth Low Energy transceiver; the CMOS neural chip, which contains the pulse-based amplifier array; and the wireless power supply, which consists of the LC resonant circuit made by the power coil and *C*_*resonant*_, a bridge rectifier to convert the AC into DC, and a 3.3V regulator which supplies the SOC and CMOS neural chip.

## III. INDUCTIVE WIRELESS POWER TRANSFER LINK

The fully wireless implantable device is powered by the use of a wireless power transfer link. This link consists of two sections, the external transmitter, and the wireless receiver coil on the flexible substrate. The system is shown in Fig. 3. Briefly, the transmitter provides energy to the transmit coil which is inductively coupled to the receive coil on the neural interface, which resonates with this energy and is rectified and regulated to a stable voltage to power the electronics on the implant.

**Fig. 3.**
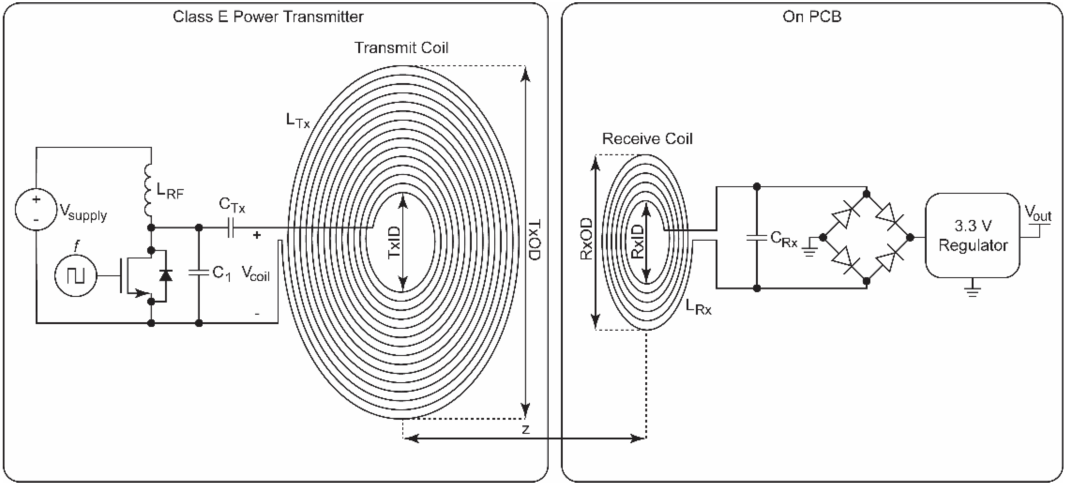
The overall structure of the wireless power transfer system. The system is based on a Class E power transmitter that drives the transmit coil at resonance, and the on PCB receive coil that is tuned to resonate at the same frequency. The optimal dimensions of the coils are determined by simulations based on previous studies [REF].

### A. Inductive link coils

The most important components of the wireless power transfer link are the transmit and receive antennas. Previously, a method has been described which optimizes the coupling coefficient between two coils for wireless power transfer by carefully selecting their dimensions [14]. The method provides detailed data to choose the optimum dimensions based on two of the coil parameters presented in Fig. 3, *z* (the distance between the two coils) and *RxOD* (the receive coil’s outer diameter). For this neural interface, the coil separation *z* was set to 15mm as it is sufficiently large to fit the implant beneath the skull and skin. *RxOD* was also chosen to be 15 mm to be reasonably small and minimally invasive while still providing sufficient space for the electronics required for the implant’s operation. The other coil dimensions, *RxID* (the receive coil’s inner diameter), *TxOD* (the transmit coil’s outer diameter), and *TxID* (the transmit coil’s inner diameter) were chosen based on HFSS simulations.

### B. Class E amplifier

For the most efficient operation of the wireless power link, the transmit coil is driven by a class E amplifier. The schematic for the class E amplifier is shown on the left side of Fig.3. The amplifier is based on a single power MOSFET that is driven by a logic-level square wave at the desired operating frequency *f. L_RF_* is an RF choke which is made sufficiently large for *f. C*_*Tx*_ is set so that a resonance occurs with *L*_*Tx*_ at *f*. The component values are chosen to maximize power delivery [19]. Power delivery to the transmit coil can be adjusted by varying *V*_*supply*_.

## IV. NEURAL AMPLIFIERS AND MODULATION CIRCUITRY

This section describes the circuitry used within the CMOS neural array chip. First, the circuitry of the pulse-based amplifiers is described, followed by a theory of operation of the pulse-based modulation technique.

### A. Neural amplifier circuitry

The schematic for the enhanced delta modulator is shown in Fig. 4. The circuit is two main parts, the amplifier section and the comparator section. The amplifier section consists of two operational amplifiers (OPAs) which are used as inverting amplifiers with capacitors as the feedback path [20]. To keep the OPAs stable at DC, high-value resistors are used in addition to the capacitors. The resistors are implemented using PMOS transistors connected as shown where they act as pseudoresistors, rather than traditional poly resistors which would occupy considerably more die area for a given resistance. This allows the output of each OPA stage to be approximately *V*_*pos*_ with no signal applied to the circuit. *V*_*pos*_ is generated in an on-chip voltage regulator and is set to ∼1.5 V, to put the DC operating point of the circuit near the middle of the amplifiers’ dynamic range. The OPAs used in this design are a cascode differential amplifier with PMOS inputs [21]–[23]. Using capacitors as the feedback elements allows for the amplifier’s gain to be independent of frequency. Examining the case of the first stage at the OPA’s inverting input:

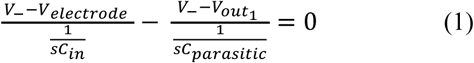

**Fig. 4.**
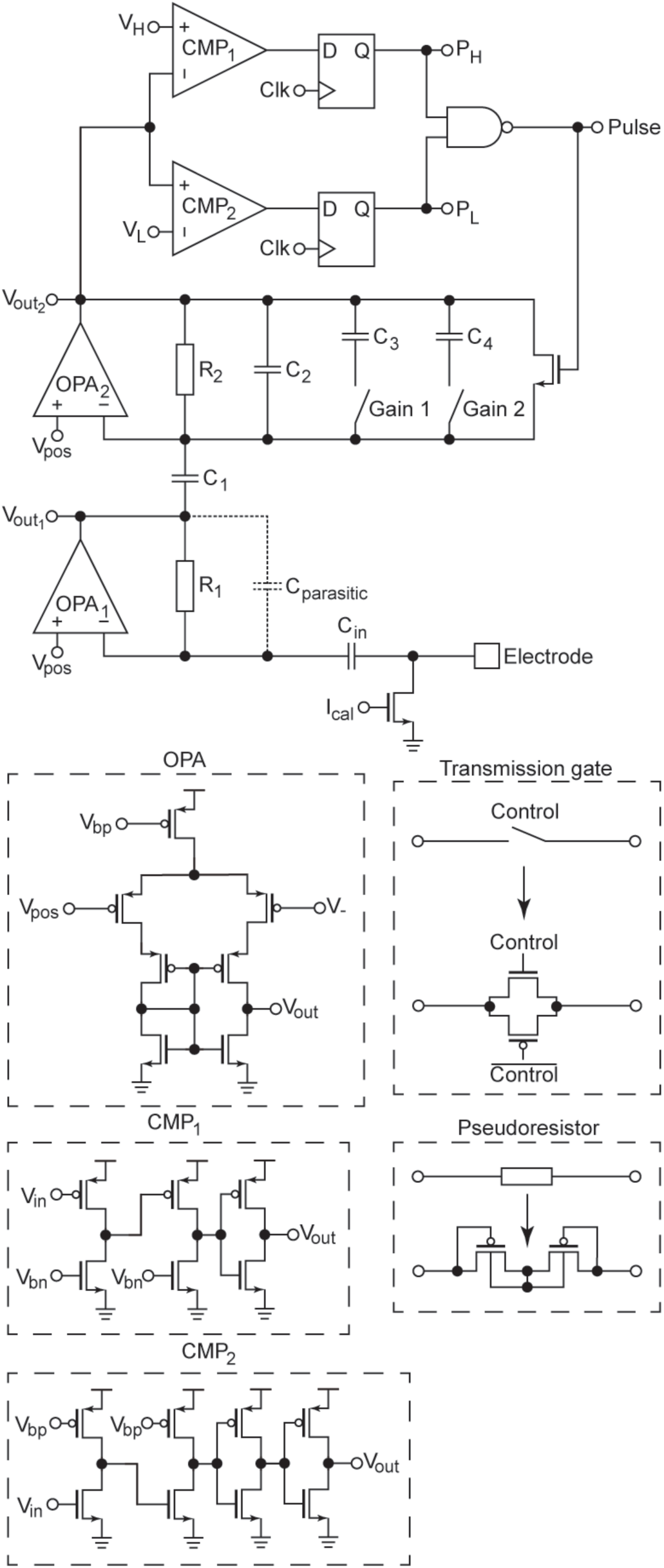
Schematic of the enhanced delta modulator neural amplifier. The amplifier uses two operational amplifiers (OPAs) with capacitive feedback to produce a frequency-independent gain. The second stage can be set to multiple gain settings using transmission gate switches to add additional capacitance to the feedback path. Two comparators are used to discharge the feedback capacitance on the second stage in order to produce a stream of bits from PH and PL whose space in time correspond to the slope of the input signal. The OPAs are a simple differential amplifier with additional cascode transistors to increase bandwidth. The comparators are a simple cascaded common-source amplifiers followed by inverters to square up the pulses. These pulses are latched by the clock signal and combined by the NAND gate to drive the reset transistor.

Notice that because *R*_*1*_’s resistance is extremely high, typically in the range of TΩ, it can be ignored. Rearranging Eq. (1):

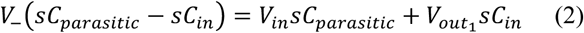

The feedback of the OPA maintains *V*_*-*_ = *V*_*pos*_. Since *V*_*pos*_ is a constant DC value, it is simply 0 in this AC analysis:

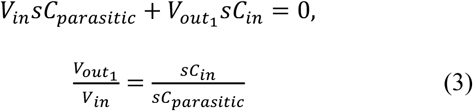

It is clear from Eq. (3) that the Laplace variable *s* can be cancelled from the expression and the gain is simply:

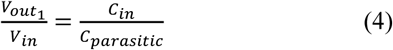

Thus, the gain of the amplifier does not depend on the frequency of the input signal. It is also only dependent on the values of the capacitors used. In the first stage, the value of *C*_*in*_ is made as large as the layout allows (∼ 1’s of pF) and the value of *C*_*parasitic*_ is determined only by the capacitance between the output and inverting input of the OPA caused by the parasitic capacitance of the transistors. The first stage, therefore, has a high gain. The second stage uses a fixed value of capacitance for *C*_*1*_ and *C*_*2*_, but has the flexibility of adding additional capacitance (*C*_*3*_ and *C*_*4*_) in parallel with *C*_*2*_ to reduce the amplifier’s gain. This is accomplished through the use of CMOS transmission gates that can be controlled via input pins to the chip.

The output of the second stage 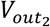 serves as the input to the comparator section of the pulse modulator. These comparators are based on two cascaded stages of common-source amplifiers for a high gain, with a PMOS input for the high threshold (*V*_*H*_) detection and an NMOS input for the low threshold (*V*_*L*_) detection. *V*_*H*_ and *V*_*L*_ are determined by the biasing level of the common-source amplifiers, set by *V*_*bp*_ and *V*_*bn*_. The outputs of the comparators are followed by an inverter (for CMP1) or two inverters (for CMP2) in order to guarantee the pulses are properly digitized and of the correct polarity. The comparators are designed with the simple cascaded common-source construction to reduce die area when compared to a larger differential comparator.

The outputs of the comparators are fed into clocked D flip-flops to allow for synchronous operation with the chip’s global clock (which enables the array’s multiplexed readout of each individual DM) and to guarantee that each reset pulse is of sufficient length to fully discharge the feedback capacitors. The two pulses, *P*_*H*_ and *P*_*L*_, are combined via the NAND gate and fed to the reset MOSFET that discharges the feedback capacitors. These pulses result in the enhanced delta modulation of the input signal which will be explained in the following section.

### B. Pulse-based neural amplifier

The pulse-based neural amplifier design allows the operation of the presented 1024 channel neural amplifier array without the need of any ADCs. The pulse-based modulation used is a type of analog-to-digital conversion that encodes the change in a signal’s voltage within the distance between digital pulses (hence delta) rather than a binary-coded representation of the analog value versus time (sampling) A conceptual study shown in Fig. 5 is performed using a MATLAB simulation. An input signal,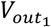, represents the output of the first stage of amplification. As 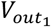 entersthe second stage, additional gain is applied to this signal, representing the gain produced by the second stage of amplification. If no delta modulation was being performed, no resets would occur and 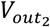 would simply be a larger amplitude (albeit with inverted sign) replica of 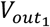. However, since modulation is occurring, once 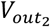 reaches either of the thresholds (*V*_*L*_ or *V*_*H*_), the corresponding pulse (*P*_*L*_ or *P*_*H*_) is triggered, and 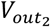 is reset to 0. In this example, as the slope of the input is rising until *t* ≈ 250, only low-going pulses (*P*_*L*_) are triggered. As the sine wave reaches its peak, and therefore its derivative (delta) approaches zero, the distance between pulses increases. This is the means by which the input signal becomes modulated.

**Fig. 5.**
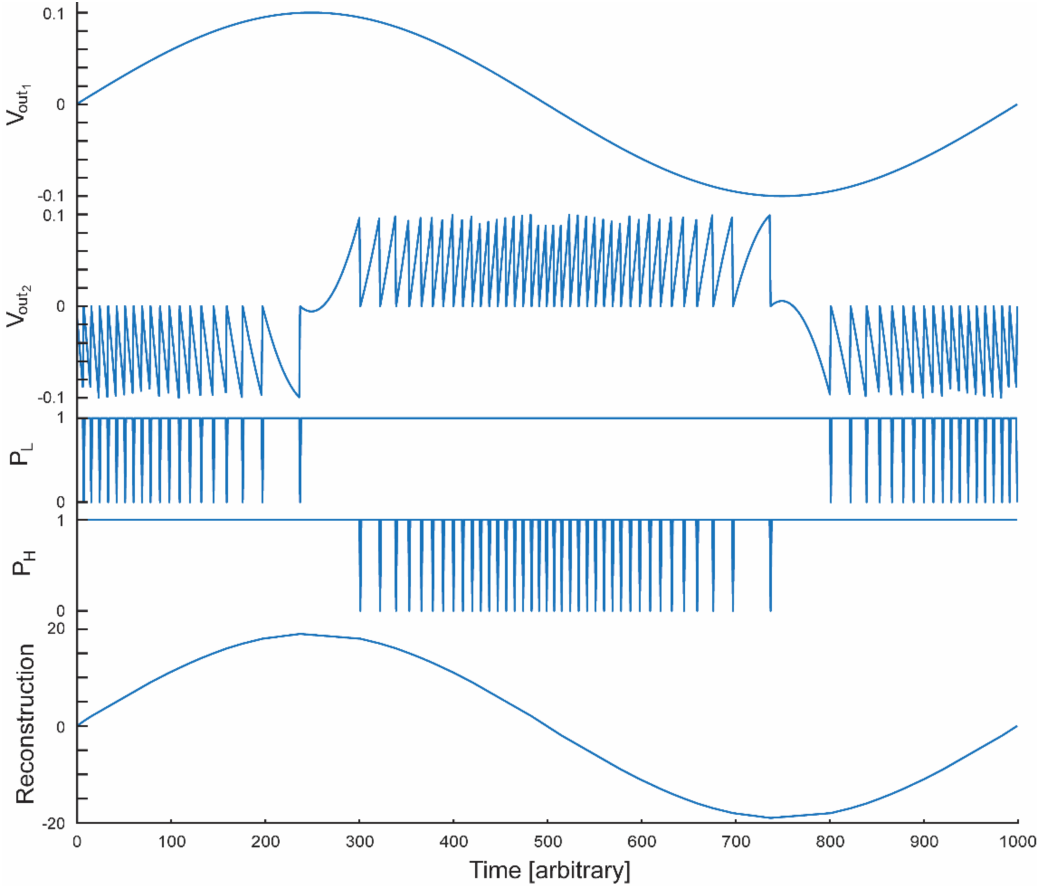
Simulation results demonstrating the enhanced delta modulation of a sine wave and the reconstruction of the signal. V_out1_ shows the input sine wave. V_out2_ shows the output of the second stage, showing the effect of the reset pulses. P_L_ and P_H_ are the pulses resulting from the comparators triggering once V_out2_ reaches either threshold V_L_ or V_H_ respectively. Reconstruction is the resulting analog reconstruction of the pulses from P_L_ and P_H_. The time dimension is unitless.

The pulses generated from the neural amplifier contains comprehensive information to demodulate and reconstruct the original signal. The analog signal is reconstructed by stepping through the pulse information. Once a pulse is detected (in this case, a zero value in Fig. 5), the output value is incremented by one unit if the pulse is from *P*_*L*_, and decremented if the pulse is from *P*_*H*_. Additionally, the output value is interpolated between pulses to avoid “stair-stepping” of the output caused by the discrete nature of the pulses. In this example, the resulting waveform is shown as “Reconstruction” in Fig. 5. The units for the reconstruction are number of pulses, if the amplitude information is important, the reconstructed value must be scaled appropriately based on knowledge of the amplifier’s gain and the threshold voltages. For the purpose of this example, the amplitude of the input wave is 0.1, the gain of the amplifier is 20, and the threshold voltages are ± 0.1. Thus, the threshold will be triggered 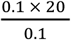, or 20 times and the reconstructed wave’s amplitude is 20 as expected. The resulting reconstructed wave in this example shows how effective the pulse-based method of modulation is at encoding and decoding an analog signal. Pulse-based modulation’s greatest weakness is obvious around *t* ≈ 250 and *t* ≈ 750, where the time between pulses increases and vertical resolution is lost, resulting in sharper transitions in the output waveform.

## V. RESULTS

The flexible neural interface device is shown in Fig. 6. The polyimide substrate is easily flexible and can be bent with the fingertips (Fig. 6a.) The size of the flexible substrate is also evident when compared with the fingertips. Fig. 6b. shows the top side of the neural interface with the SOC (towards the top), the BLE antenna (the white object in the middle), and the voltage regulator (small square IC towards the bottom and right) easily visible. Fig. 6c. shows the bottom side of the interface. The CMOS chip, integrating the neural amplifier array, is attached to the back side of the flexible neural interface. The wireless power transfer coil is also positioned on the back side around the perimeter. In this section, the performance of the wireless power transfer system, the pulse-based modulators on the CMOS chip, and the amplifier array is discussed.

**Fig. 6.**
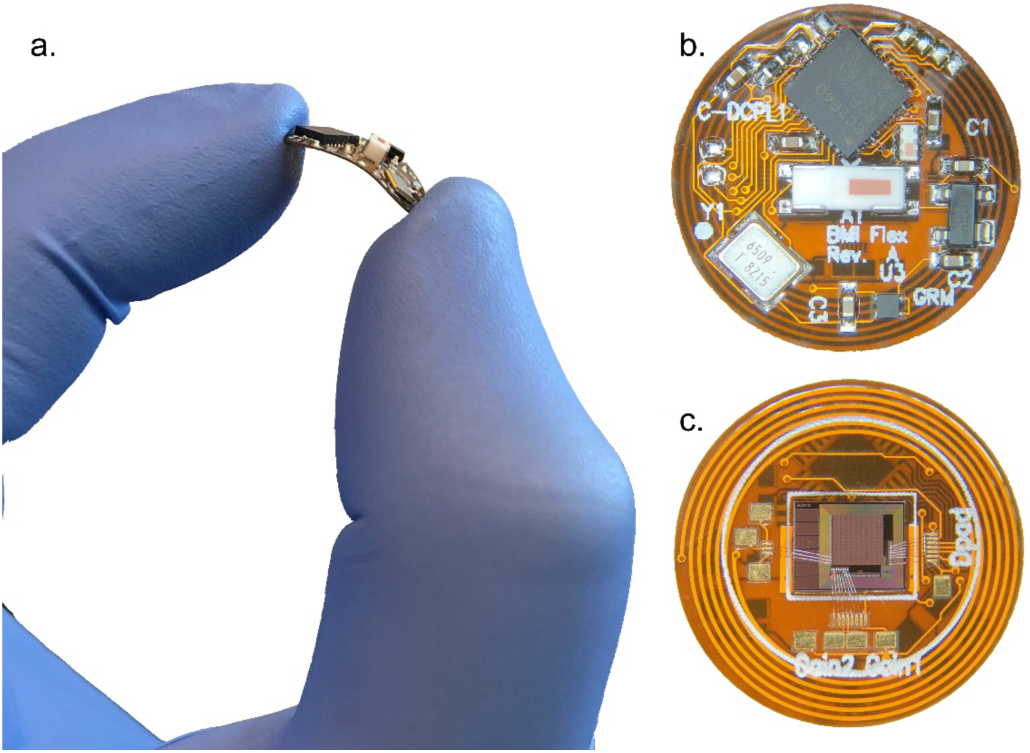
Photos of the implantable neural interface. a. the neural interface being flexed by a thumb and forefinger. b. the top side of the neural interface. c. the bottom side of the neural interface.

### A. Wireless power transfer performance

The wireless power transfer performance is characterized by monitoring the regulated output while altering the operating frequency, transmitter power, as well as distance between the inductive link coils. The wireless power transfer characteristics of the system are shown in Fig. 7. Fig. 7a. displays the regulated output voltage across a range of input frequencies from 2.3 to 2.9 MHz. The output plateaus around the chosen desired operating frequency of *f* = 2.64 MHz due to the effect of the voltage regulator. Fig. 7b. shows the voltage across the transmit coil across the same range of operating frequencies. The class E amplifier is tuned for resonance near the desired operating frequency of 2.64 MHz, so its performance drops when operating away from the desired operating frequency. Fig. 7c. shows the regulated output voltage versus the supply voltage to the class E transmitter varied from 0 to 15 V at a fixed frequency of *f* = 2.64 MHz and distance *z* = 15 mm. This shows clearly the effect of voltage regulator once the supply voltage reaches 9 volts. At this point there is sufficient power being delivered that the regulator is able to supply a constant output voltage. Fig. 7d. shows the regulated output voltage with respect to the distance *z* between the transmit and receive antennas being swept from 10 to 29 mm. The effect of the voltage regulator can again be seen by keeping the output voltage stable below *z* = 15 mm, which is the designed nominal working distance. These results show that the wireless power transfer system is functioning as designed and expected.

**Fig. 7.**
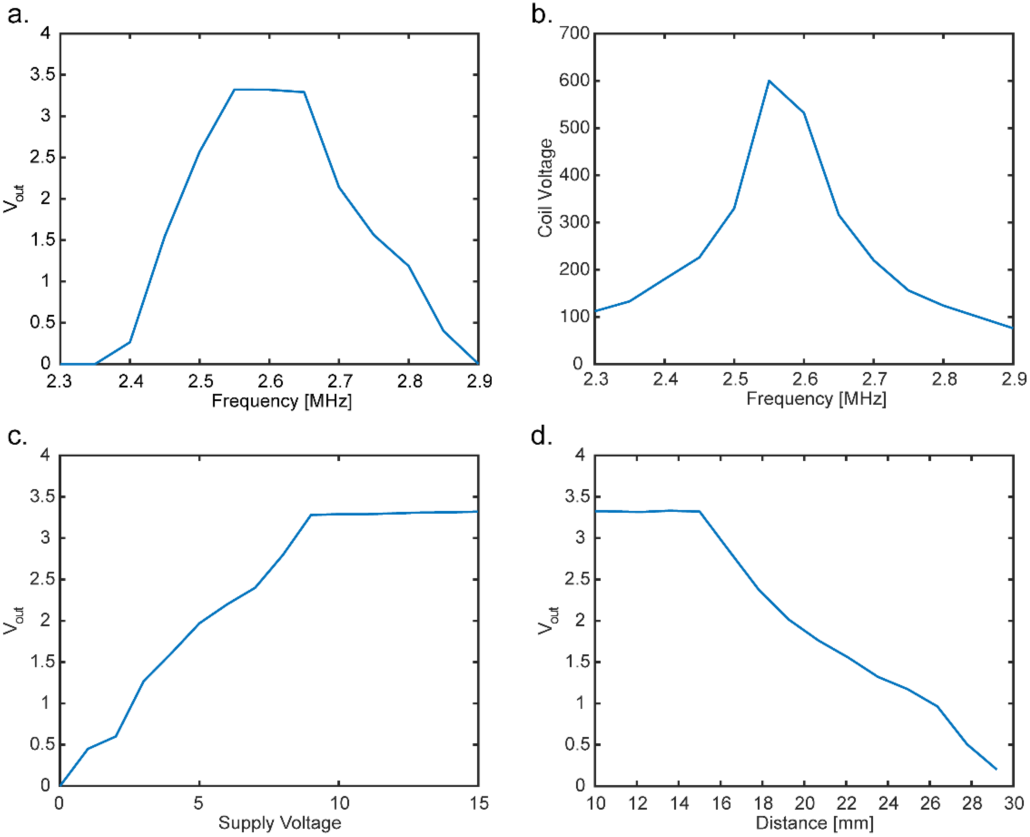
Plots showing the performance of the wireless power supply. a. regulated output versus the power frequency, the output is a stable 3.3 V around the desired 2.64 MHz nominal operating frequency. b. the power transmit coil’s voltage versus frequency, the highest output voltage peak is caused by the transmitter’s resonant nature. c. the regulated output voltage versus the voltage applied to the class E power transmitter, above ∼9 V the output is stabilized at 3.3 V. d. the regulated output voltage versus the distance between the power transmit coil and the implant’s receive coil, performance decays after the designed working distance of 15 mm.

### B. Pulse-based modulation / demodulation on the CMOS chip

The CMOS neural interface chip embeds an array of 1024 pulse-based neural amplifiers. Additionally, it also contains a single testing amplifier/modulator that can be operated independently from other amplifiers and signals that are not normally accessible such as the output of the first amplification stage (*V*_*out1*_), the second amplification stage (*V*_*out2*_), and the digital pulses out (*P*_*L*_ and *P*_*H*_) can be measured. This “testing amplifier” utilizes the exact same circuitry as the rest of the 1024 arrayed amplifiers, so the additional taps allow for a visualization of what is occurring within each amplifier in the array.

The testing amplifier is stimulated with a large amplitude (7 mV) sine wave, with the clock running at 50 kHz. We selected the large amplitude to better visualize the pulses and resets occurring during measurements in this particular experiment. The results of this experiment showing waveforms from the *V*_*out1*_, *V*_*out2*_, *P*_*L*_, *and P*_*H*_ nodes are displayed in Fig. 8. The reconstructed signal is produced using the same method described in section IV. B. (delta modulation). Although this experiment allows for a detailed interrogation of the neural amplifier at different nodes, it is worth noting that the testing circuit does not accurately match the performance of amplifiers within the array. The additional connections to sensitive nodes within the amplifier adds parasitic capacitance at each node and affects the characteristics such as gain, noise and bandwidth. Thus, the results of Fig. 8, aren’t a completely accurate representation of what is occurring within amplifiers in the array. However, it does offer an excellent visual aid to understand the pulse-based amplifier’s operation. The input signal is first amplified and shown at *V*_*out1*_. The signal is passed to the second stage amplifier, where the comparators trigger the reset pulses, causing *V*_*out2*_ to fall back to zero and the appropriate high or low pulse to be output. Because of the comparators’ simple design, the comparators do not fire at consistent delta sizes. This effect is obvious when looking at the difference in the number of low-going pulses versus high-going pulses in Fig. 8 (*P*_*L*_ and *P*_*H*_). Because of this asymmetry in delta, an additional step of processing is needed to produce the reconstructed result. Although the comparator thresholds are not consistent with each other, they are consistent with themselves. This has the result of adding a constant slope to the reconstructed output, which is easily subtracted, producing the reconstruction as expected. The resulting reconstruction of this sinusoidal input is shown to follow the input as expected.

**Fig. 8.**
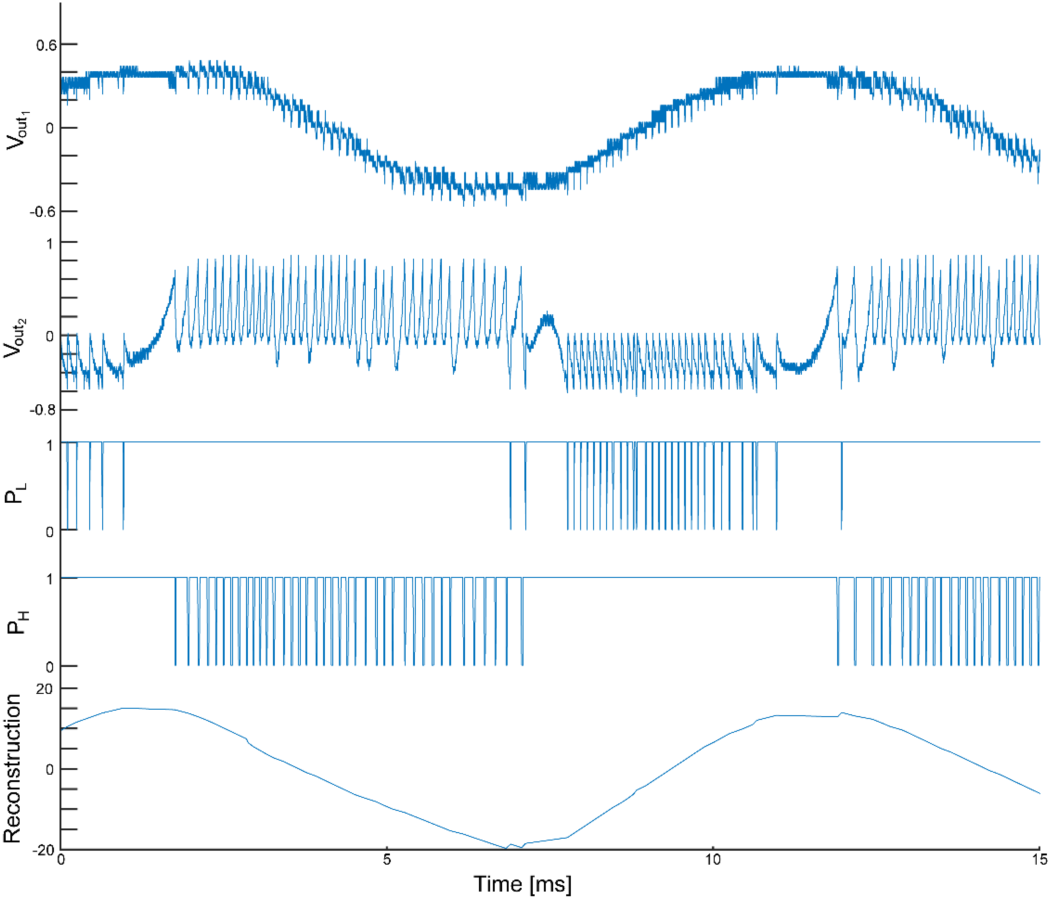
Measured results from the testing amplifier. V_out1_ shows the output of the first stage of amplification. V_out2_ shows the output of the second stage. P_L_ and P_H_ are the digital pulses from the comparators. Reconstruction is the resulting reconstruction using the pulses

### C. Array data via bluetooth

To demonstrate the functionality of the complete system with wireless power transfer and wireless data transmission, a flexible neural interface was assembled with one of these wire-bonded pads connected to a signal generator. A 5 mV amplitude sine wave at 1 kHz was injected into the electrode. The neural interface was also positioned 15 mm from the class E amplifier’s transmit coil, which was powered by 10 V. The data was streamed out of the neural interface via BLE to a computer and was processed in MATLAB to yield the FFT shown in Fig. 9. A large peak is visible above the noise directly at the 1 kHz applied. Because of the poor performance of the comparators, however, the 1/f shape to the noise floor is clearly visible as well. Since this noise is so large, this data is presented in the frequency domain rather than the time domain, as the noise obscures the visibility of the 1 kHz signal from the human eye in the time domain. Although a time domain signal is preferred, FFTs can be used to extract information from EEG signals [24]. However, with the current performance of the comparators, signals below ∼1 kHz are not currently detectable due to the 1/f noise.

**Fig. 9.**
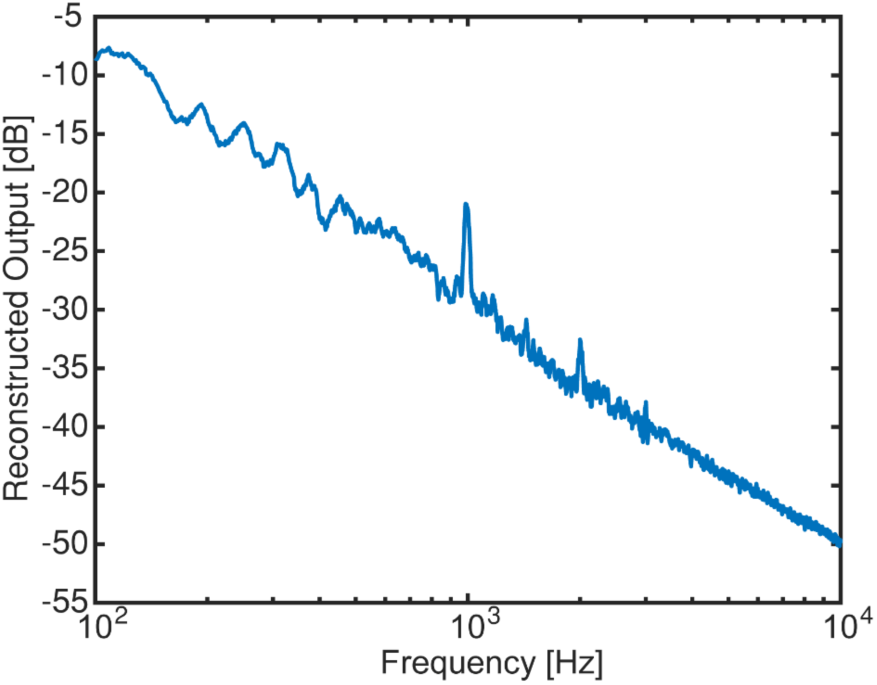
FFT spectrum of a reconstructed output from the CMOS neural chip. The input signal was a 5 mV peak 1 kHz sine wave. The implant was wirelessly powered for this measurement.

## VI. CONCLUSION

A compact, flexible neural recording interface has been developed and tested to operate on wireless power and with wireless data transmission. Such a device is capable, with the addition of passivating materials such as PDMS, of being permanently implantable in a human to continuously study the brain without the need of transcranial connectors. Work is actively being done to improve the design of the implant. The CMOS chip’s current largest weakness is the design of the comparators as can be seen by the results. Since they were designed to be as compact as possible, their performance, mainly 1/f noise, has suffered as a result. A similar CMOS device with improved comparators will increase performance. Other improvements to the system will enable the simultaneous readout of all 1024 neural recording amplifiers by using a higher bandwidth data transfer link, and further reduce the size of the implant by integrating more of the electronics onto the chip. However, despite the current limitations of the device, it has still proven possible to operate on wireless power, amplify and modulate signals using enhanced delta modulation, and wirelessly transmit data.

